# Latch Verified Bulk-RNA Seq toolkit: a cloud-based suite of workflows for bulk RNA-seq quality control, analysis, and functional enrichment

**DOI:** 10.1101/2022.11.10.516016

**Authors:** Hannah G.B.H. Le, Jacob L. Steenwyk, Nathan Manske, Max Smolin, Aidan Abdulali, Ayush Kamat, Rohan Kanchana, Kyle Giffin, Alfredo Andere, Kenny Workman

**Author notes:** These authors contributed equally (Hannah G.B.H. Le, Jacob L. Steenwyk). Correspondence should be addressed to Hannah G.B.H. Le or Kenny Workman.

## Abstract

**Background:** Analysis of high-throughput bulk RNA-sequencing (RNA-seq) data reveals changes in gene expression between diverse conditions. Many tools have emerged to quality control RNA-seq reads, quantify expression levels, conduct functional enrichment among differentially expressed genes, or identify differential RNA splicing. However, unified toolkits for conducting these analyses are lacking. Moreover, existing software does not use cloud-based platforms that provide the necessary storage and computational resources to process RNA-seq data or intuitive graphical interfaces for easy use by experimental and computational scientists.

**Results:** To address these challenges, we introduce the Latch Verified Bulk RNA-Seq (LVBRS) toolkit, a flexible suite of programs packaged into a single workflow coupled with a graphical user interface for conducting quality control, transcript quantification, differential splicing, differential expression analysis, and functional enrichment analyses. For functional enrichment, the LVBRS toolkit supports three databases—Gene Ontology, KEGG Pathway, and Molecular Signatures database—capturing diverse functional information. We demonstrate the utility of the LVBRS toolkit by reanalyzing a publicly available dataset examining the impact of severe and mild models of hypoxia—induced by Cobalt (II) Chloride (CoCl_2_) and oxyquinoline treatment, respectively—on a human colon adenocarcinoma cell line. Our analyses reveal CoCl_2_ treatment results in more differentially expressed genes, recapitulating previously reported results that CoCl_2_ models more severe hypoxia. Moreover, including alternative splicing and functional enrichment analysis using a greater breadth of functional databases revealed additional biological insights—such as greater alternative splicing in the CoCl_2_ condition and differentially expressed DNA repair pathways. These results demonstrate the LVBRS toolkit’s efficacy in facilitating biological insights from bulk RNA-seq data.

**Conclusions:** The LVBRS toolkit offers a robust unified framework for processing and analyzing Bulk RNA-Seq experiments. The easy-to-use graphical user interface will enable diverse scientists to conduct high-throughput bulk RNA-Seq analysis efficiently. Our aim is that the LVBRS toolkit will help streamline bulk RNA-seq workflows and facilitate deriving biologically meaningful insights from bulk RNA-seq data. The source code is freely available under the MIT license and hosted on the LatchBio Console (https://console.latch.bio/se/bulk-rnaseq), complete with documentation (https://latch.wiki/bulk-rna-seq-end-to-end).

## Background

Next-generation sequencing has deepened our understanding of genomes, transforming numerous biological disciplines [1]. Several technologies exist to probe different aspects of genome organization and function. For example, chromatin immunoprecipitation followed by sequencing (ChIP-seq) can facilitate mapping genome-wide profiles of DNA-binding proteins and histone modifications [2]. One technique that has become prominent is bulk RNA-sequencing (RNA-Seq).

Bulk RNA-seq has become a ubiquitous tool for gaining insight into the transcriptomes of diverse organisms [3]. Advances have enabled isoform discovery, *de novo* transcriptome analysis, fusion transcript discovery, structural and RNA–protein interaction analysis, and more [4]. A common use of bulk RNA-seq is differential gene expression (DGE) analysis, a method to determine the quantitative changes in expression levels between a control condition and experimental treatments using short-read sequencing technologies [5]. Similarly, recent developments have enabled the analysis of differential splicing between control and experimental groups [6]. Thereafter, pathway enrichment analysis is often conducted, facilitating insights into biological pathways with modified expression or splicing [7].

Despite the widespread use of bulk RNA-seq, DGE, alternative splicing, and enrichment analysis, there are few automated high-throughput tools to analyze bulk RNA-seq data and conduct pathway enrichment analysis [8–11]. For example, the tool RAP provides a cloud-based service for bulk RNA-seq quality control, transcript count matrix generation, and DGE analysis but no pathway enrichment analysis [9]. For RAP, users are restricted to two projects, twelve FASTQ files, and a 30-day file storage policy, stymying adoption for large-scale and long-term projects. Another tool, WEBGIVI, facilitates gene enrichment analysis and visualization but does not handle RNA-seq data [10]. Notwithstanding the utility of RAP and WEBGIVI, the lack of a unified toolkit may be due to a host of challenges, including difficult-to-maintain workflows, requiring extensive and heterogeneous compute resources, demanding proficiency in cloud infrastructure engineering, and file system management and performant networking, resulting in a high barrier to entry for conducting RNA-seq and DGE analysis [12, 13].

To address these issues, we developed the Latch Verified Bulk RNA-Seq (LVBRS) toolkit, a suite of end-to-end bioinformatic workflows with automated plotting of results for immediate and easy interpretation. The LVBRS toolkit is available on a cloud-based infrastructure designed to alleviate users from computational and data management burdens and streamline analysis. Reads are quality-trimmed and can be mapped to a genome available in a curated and pre-indexed set of reference genomes; alternatively, users can upload a custom genome, enabling the LVBRS toolkit to be used on organisms from across the tree of life. DGE, differential splicing, and functional enrichment analysis place results in a broader biological context. The LVBRS toolkit is open-source; thus, users can modify steps as needed using the Latch software development kit (https://docs.latch.bio/) and still benefit from a cloud-based infrastructure. To demonstrate the utility of this workflow, we reanalyzed publicly available RNA-Seq data examining the impact of severe and mild hypoxia induced by Cobalt (II) Chloride (CoCl_2_) and oxyquinoline (Oxy) treatment, respectively, on a human colon adenocarcinoma cell line [14]. LVBRS successfully recapitulated major findings. Incorporating additional functional databases and alternative splicing analysis facilitated new biological insights; for example, CoCl_2_-treatment resulted in more differentially spliced genes than Oxy-treatment, corroborating previous findings that CoCl_2_ induces more severe hypoxia [14]. The LVBRS toolkit facilitates high-throughput analysis of Bulk RNA-Seq analysis and alleviates researchers from the burden and cost of data management, workflow maintenance, and resource allocation.

### Implementation

The LVBRS toolkit features three modular workflows, enabling users to implement workflow modules or the entirety thereof (Figure 1). One module is for quality trimming, alignment, gene-wise transcript quantification, and differential intron excision analysis, complete with summary information about RNA-Seq read quality. The second module is for DGE analysis between experimental groups. The third module is for functional enrichment analysis. The modular design of the LVBRS toolkit accommodates diverse use cases and desired outputs from users.

**Figure 1.**
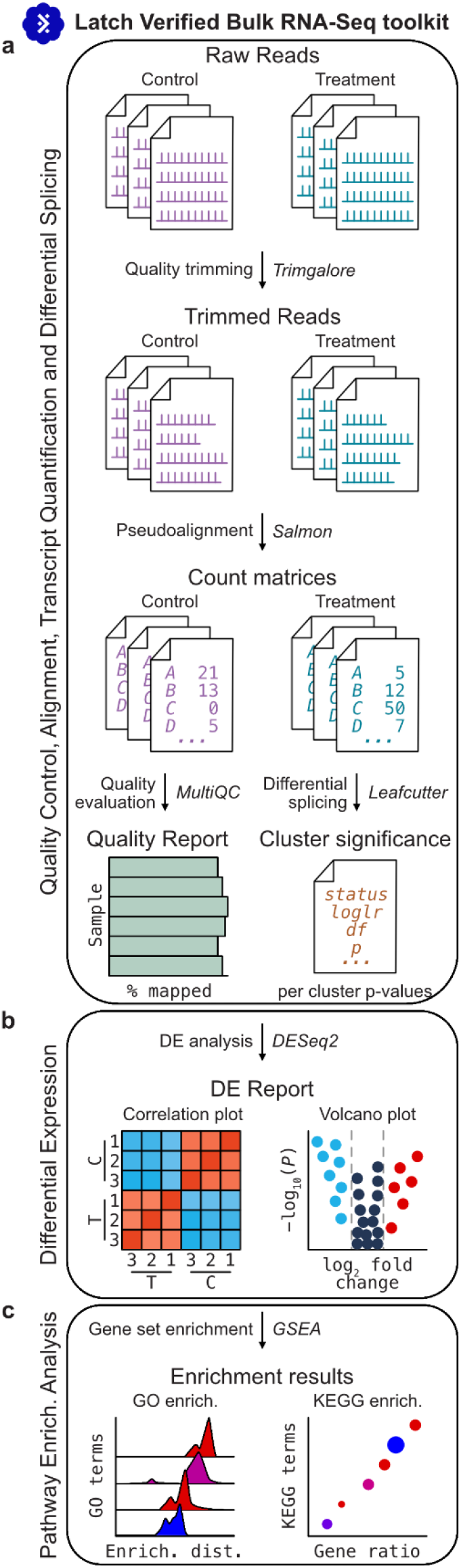
The LVBRS toolkit for RNA-sequencing quality control, mapping, and enrichment analysis. The LVBRS toolkit is composed of three workflows, allowing flexible user implementation. (a) The first workflow, Bulk RNA-seq, conducts quality trimming and aligns the resulting reads to generate count matrices. Quality reports detailing read lengths and the number and percentages of mapped reads are also provided. An optional toggle can be switched on to conduct alternative splicing analysis. (b) A workflow for differential expression analysis also generates interactive correlation plots across all samples and volcano plots. (c) The pathway enrichment analysis workflow conducts functional enrichment analysis using three ontology databases: GO, KEGG, and MSigDB. Outputs from each step are depicted as cartoons here and are not exhaustive representations of the output generated by the LVBRS.

### Bulk RNA-Seq quality control, alignment, transcript quantification, and alternative splicing detection

The LVBRS workflow for quality control, alignment, and transcript quantification ingests short-read sequencing files in FASTQ format. Sequence reads are quality-trimmed using TrimGalore, v0.6.6 (https://www.bioinformatics.babraham.ac.uk/projects/trim_galore), using default parameters. The resulting quality-trimmed reads are mapped to a reference genome using Salmon, v1.8.0 [15], with default parameters. Reads are mapped against a genome from a precomputed database of several organisms—*Homo sapiens, Mus musculus*, and *Saccharomyces cerevisiae* (Table 1) [16–19]—or a custom reference genome in FASTA format, along with an annotation file in gene transfer format (gtf) to describe gene boundaries. The LVBRS toolkit uses FastQC (http://www.bioinformatics.babraham.ac.uk/projects/fastqc/) to generate a quality control report of trimmed FASTQ files. Also, a MultiQC report is generated to detail mapping results—the number and percentage of reads mapped per sample and fragment length distributions [20]. A count matrix representing the number of reads mapped per gene is generated from the Salmon results.

**Table 1.**
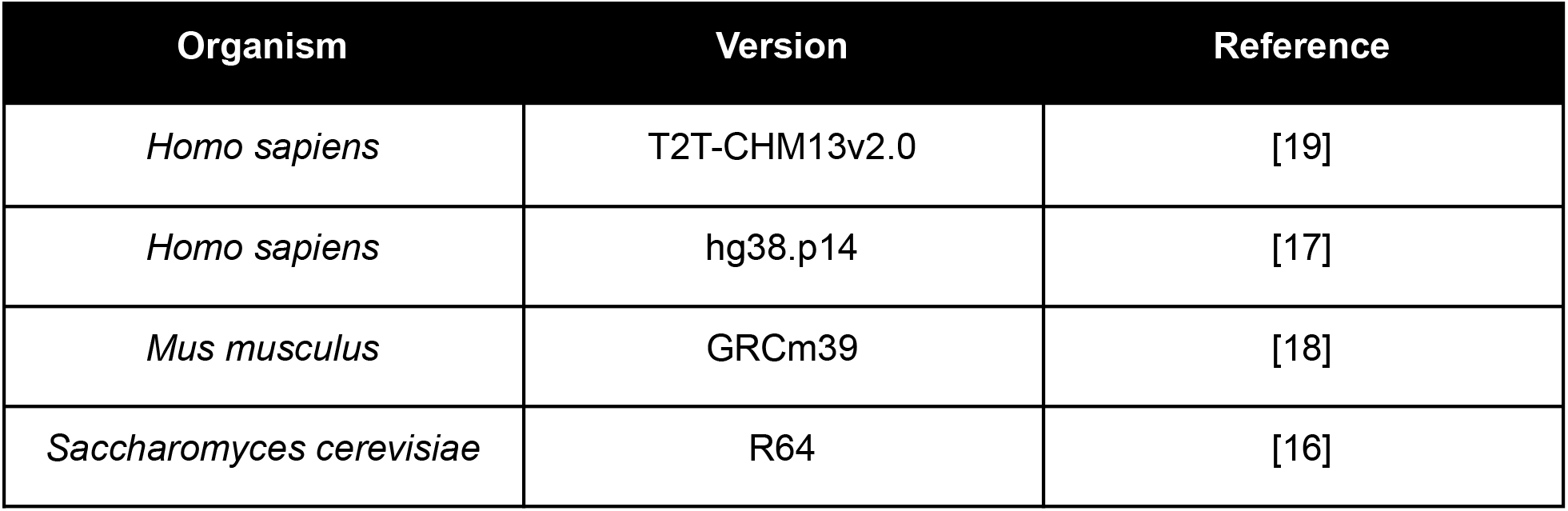
Genomes in the Latch Genome Database. The Latch Genome Database comes complete with four commonly used reference genomes. The Latch Genome Database is an open-source database (https://github.com/latchbio/latch-genomes).

| Organism | Version | Reference |
| --- | --- | --- |
| *Homo sapiens* | T2T-CHM13v2.0 | [19] |
| *Homo sapiens* | hg38.p14 | [17] |
| *Mus musculus* | GRCm39 | [18] |
| *Saccharomyces cerevisiae* | R64 | [16] |

The LVBRS workflow also conducts differential intron excision analysis (also known as differential splicing analysis) between two conditions using LeafCutter, v0.2.9 [6]. To include LeafCutter in workflow execution, users click a toggle button from off to on. In doing so, three steps are automatically included in the workflow execution. In the first step, the outputted alignment files in bam format, which were generated by Salmon [15], are converted to junction files [21]. In the second step, the junction files are used to cluster together introns; a minimum of 50 split reads supporting each cluster is required, and introns of up to 500 kilobases are allowed. In the third step, differential intron excision analysis is performed, generating summary files that output significant differences between clusters and the effect sizes per intron in the conditions. The three steps for differential intron excision analysis follow best practices described in the LeafCutter documentation [6].

In summary, this workflow takes a minimum input of reads in FASTQ format, trims low-quality reads and adapters, maps reads, outputs read counts per gene, quality reports, and conducts differential intron excision analysis.

### Differential Gene Expression Analysis

The LVBRS workflow for differential gene expression analysis takes as input RNA-seq transcript counts per gene, the output of the previous workflow. DESeq2, v3.15 [22], is used to identify differentially expressed genes between experimental conditions. DESeq2 was chosen because a benchmarking study found that DESeq2 had the highest true positive rate, specificity, and accuracy, among other measures of performance [23].

The LVBRS implements a visualizer server, similar to an RShiny App or DashApp, for intuitive exploration and interpretation of results. Plots can be described by three broad categories, providing a hierarchical view of results. The highest-order category, “overview,” provides a global view of the data. The next order category, “contrast,” provides pairwise comparisons between experimental groups. The lowest-order category, “genes of interest,” allows users to plot the normalized expression levels of specific genes. Among “overview” plots, a correlation matrix plot and a plot of each sample in principal component space enable users to compare global similarities and differences between samples (Figure S1 and S2). Additional summary figures—such as a heatmap of genes with the highest read counts (Figure S3) and a gene size factor distribution plot, depicting the distribution of read counts per gene, based on the geometric mean (Figure S4)—are also generated. Among “contrast” plots, a mean-difference plot (commonly referred to as an MA-plot), depicts gene-wise log fold-change compared to mean expression between two treatment groups [24]; genes that are significantly differentially expressed are noted using a color scheme (Figure S5). A volcano plot is also generated to shed light on the distribution of genes that are over- or under-expressed in the treatment group (Figure S6). For “genes of interest”, a clustering heatmap of z-scores based on normalized read counts per gene (Figure S7).

The graphical user interface is designed to facilitate data exploration. For example, plots are interactive meaning that users can drag their mouse over data points of interest to get more information about them. For example, users that hover their mouse over a data point in a volcano plot will get gene name, log_2_ fold change, and -log_10_(*P*) value information. Where applicable, figures often feature hierarchical clustering analysis to further highlight similarities and differences between samples. All raw data used to generate plots and portable network graphics can be downloaded.

### Gene Set Enrichment Analysis

The LVBRS workflow for functional enrichment analysis implements the Gene Set Enrichment Analysis framework [7] and takes the output of the DEG workflow as input. This workflow conducts enrichment analysis using three databases—the Gene Ontology (GO) database [25], KEGG Pathway database [26], and the Molecular Signatures Database (MSigDB) [7]—capturing diverse functional information. Nuanced differences are implemented during enrichment calculations for each database, however, the same three-step statistical framework is implemented. First, an enrichment score is calculated; second, the significance level is assessed; third, p-values are adjusted for multiple hypothesis testing. The gene set enrichment analysis method implements a Kolmogorov—Smirnov-like statistic, a permutation test to generate a null distribution for p-value determination, and a false discovery rate analysis to reduce false positives.

The LVBRS workflow generates several informative figures summarizing enrichment results. For each database, three panels are generated. Two panels are dot plots depicting gene ratio plots for gene sets with increased or decreased differential expression (Figure S8). The size of each dot reflects the count of genes associated with the pathway in the differentially expressed gene list, and the color reflects the multi-test corrected p-value. In the third panel, a ridge plot depicts a density plot of fold change values per gene within each enriched term (Figure S9). Moreover, pathway figures are also generated using KEGG information and Pathview [27] (see Figure S10 for an exemplar figure). Lastly, raw data files summarizing enrichment per term, p-values, and other data are provided as comma-separated text files.

In summary, the LVBRS workflow for gene set enrichment analysis facilitates analysis of broadly impacted gene functions and pathways in an experimental group.

## Results

Since releasing the LVBRS toolkit to the public approximately three months ago, the workflows have been executed 4,731 times (as of November 4th, 2022). The average runtime for quality control, read mapping, transcript quantification, and summary statistics is 26 minutes and 25 seconds; for DGE analysis, the average runtime is 4 minutes and 39 seconds; and for functional enrichment analysis, the average runtime is 6 minutes and 15 seconds. The number of workflow executions demonstrates that the LVBRS toolkit can handle and process large amounts of data and a broad user base. Importantly, these results indicate users can reliably obtain biologically meaningful results in under an hour.

To demonstrate the utility of the LVBRS toolkit, we reanalyzed Bulk RNA-Seq data generated to evaluate the impact of severe and mild hypoxia—induced by the treatment of Cobalt (II) Chloride (CoCl_2_) and oxyquinoline (Oxy), respectively—on undifferentiated Caco-2 Cells [14], a human colon adenocarcinoma cell line [28]. After quality control trimming, an average and standard deviation of 89.51 ± 1.24% of reads were mapped, equating to 20.5 ± 2.03 million reads. DGE analysis identified 3,717 and 1,442 up-regulated and 3,627 and 621 down-regulated genes in CoCl_2_ and Oxy treatment, respectively (Table 2). Differential splicing analysis revealed differential intron excision between 145 and 557 clusters of introns between the Oxy treatment and control groups and the CoCl_2_ treatment and control groups, respectively (p < 0.05 after false discovery rate multi-test correction; log-likelihood ratio test). CoCl_2_ treatment resulted in a greater number of differentially expressed and spliced genes, in line with the expectation that CoCl_2_-compared to Oxy-treatment, results in more severe hypoxia.

**Table 2.** The number of up- and down-regulated genes in CoCl_2_- and Oxy-treated Caco-2 cells. Following previously established protocol [14], differentially expressed genes are defined as having a log_2_ fold change of 1 or greater and a false discovery rate of less than 0.05. Differentially expressed genes were compared to the control group, which was not treated with CoCl_2_ or Oxy. As expected, the number of differentially expressed genes is greater when Caco-2 cells are treated with CoCl_2_ than Oxy. Although these two comparisons are shown here, the LVBRS automatically conducts all pairwise comparisons.

| Treatment group | Number of up-regulated genes | Number of down-regulated genes |
| --- | --- | --- |
| CoCl<sub>2</sub> | 3,717 | 3,627 |
| Oxy | 1,442 | 621 |

Examination of DEGs and functional enrichment analysis revealed the LVBRS toolkit successfully recapitulated previously reported results [14]. Specifically, most DEGs in each study are the same (Table 3). Furthermore, enriched functional terms were the same or similar—for example, the MSig term “HALLMARK_TNFA_SIGNALING_VIA_NFKB” and terms associated with apoptosis (Figure S11). As previously reported, these results may indicate a complex interplay between hypoxia response, inflammation, and apoptosis [14]. The previously published study did not conduct enrichment analysis with the GO database; however, GO enrichment analysis conducted by the LVBRS toolkit revealed enrichment of terms associated with inflammation. Specifically, GO-enriched terms among up-regulated genes include “positive regulation of immune effector process,” “positive regulation of immune response,” and “T cell activation” (Figure S8). These results support the hypothesis that inflammation is part of the response to hypoxia and underscores the utility of incorporating functional enrichment analysis across multiple databases.

**Table 3.** Comparison of published DEGs and DEGs identified using the LVBRS toolkit. The number of DEGs identified by a previously published study [14] and the LVBRS toolkit is shown here. Overall, the majority of DEGs are the same in each study.

| Treatment group | DEGs identified in both the previously published study [14] and the LVBRs toolkit | Unique DEGs in a previously published study [14] | Unique DEGs identified using the LVBRs toolkit |
| --- | --- | --- | --- |
| CoCl<sub>2</sub> | 4,915 | 515 | 2,429 |
| Oxy | 1,422 | 205 | 641 |

The LVBRS toolkit and previously published study did, at times, differ. For example, the LVBRS toolkit includes alternative splicing detection between treatment and control groups. Alternative splicing detection revealed a greater number of differentially excised introns in the CoCl_2_ treatment group compared to the Oxy treatment group. For example, the LVBRS toolkit implicated DNA repair pathways—specifically, the KEGG terms “base excision repair” and “Fanconi anemia pathway”—to be enriched among down-regulated genes in the CoCl_2_-treated group (Figure S12). Interestingly, DNA damage response genes are known to be down-regulated in colorectal cancer cells, which can be exploited for cancer therapies [29, 30]. In other words, the LVBRS toolkit not only successfully identified a functional class of genes expected to be down-regulated by hypoxia in a model of colon cancer but also identified a known therapeutic target. GO term enrichment analysis revealed the term “unfolded protein binding,” which is part of a previously reported response to hypoxia in cancer [31], was enriched among up-regulated genes. These results highlight additional biologically relevant responses to hypoxia in a model human colon adenocarcinoma cell line.

Differences between previously published results [14] and those reported here are likely due to differences in workflow architecture. Differences between each study include that the LVBRS toolkit conducts read quality and adapter trimming, whereas the previous study conducted only the latter [14]. Furthermore, the previous study used STAR [32] for read-mapping; the LVBRS toolkit uses Salmon. A study evaluating the performance of different read aligners and the impact on downstream analysis revealed results could differ between Salmon and STAR using an RNA-Seq dataset of control and cold-acclimated conditions—for example, the percentage of mapped reads and significantly differentially expressed genes [33]. More broadly, these findings support observations that workflow architecture can impact results [34].

## Discussion

Here, we present the LVBRS toolkit, a user-friendly and cloud-based platform for conducting and deriving biological insights from bulk RNA-Seq experiments. The LVBRS toolkit implements best practices [35] and the latest software developments to conduct quality control of RNA-seq samples, quantify expression levels, conduct functional enrichment among differentially expressed genes, and identify differentially excised introns. Notably, the LVBRS toolkit conducts these analyses in a simple end-to-end framework, alleviating researchers from several hurdles, including data management and file handling, while providing the power of cloud-based computing.

Using the LVBRS toolkit to reanalyze publicly available data, we recapitulated previously published results [14] and added additional biological insights by incorporating additional functional databases and alternative splicing analysis. For example, both studies identified the MSig term “HALLMARK_TNFA_SIGNALING_VIA_NFKB” to be enriched among DEGs. Among additional biological insights, our analyses revealed the KEGG terms “base excision repair” and “Fanconi anemia pathway”—to be enriched among down-regulated genes in the CoCl_2_-treated group. Lastly, alternative splicing analysis revealed more differential excised introns in the CoCl_2_-treated group compared to Oxy. These results demonstrate the efficacy and utility of the LVBRS toolkit for conducting high-throughput analysis and deriving biologically meaningful data from Bulk RNA-Seq experiments.

The LVBRS toolkit will continue to undergo maintenance and development to ensure stable and reproducible performance. Future iterations will follow changing best practices for data processing and analysis. Additional developments will expand current resources and refine data visualization. For example, resources that will be expanded include additional genomes in the Latch database; future data visualization features will include leafviz, a toolkit for visualizing LeafCutter results.

## Conclusion

Analysis of Bulk-RNA Seq experiments is commonly employed; however, a cloud-based platform for high-throughput analysis and full support for data management and file handling support is lacking. We present the LVBRS toolkit, a suite of workflows to conduct end-to-end analysis of Bulk-RNA Seq experiments that address this gap, alleviating researchers from maintaining cumbersome workflows and computational infrastructure for research. Moreover, the LVBRS toolkit comes complete with an easy-to-use interface, enabling intuitive exploration of results and data.

## Availability and requirements

Project name: Latch-verified Bulk RNA-Seq toolkit

Project home page: https://latch.wiki/bulk-rna-seq-end-to-end

Operating system(s): Platform independent

Programming language: Python and R

Other requirements: NA

License: MIT

Any restrictions to use by non-academics: 100 credits are provided for free by default. There is option to purchase additional credits if required.

Users must create profiles to maintain data on the Latch platform. Users can create accounts associated with preexisting GitHub, Google, or Microsoft accounts.

## Declarations

### Ethics approval and consent to participate

Not applicable

## Consent for publication

Not applicable

## Availability of data and materials

The LVBRS codebase is publicly available. The workflow for conducting RNA-Seq quality control, mapping, transcript quantification, and alternative splicing is available here: https://github.com/latch-verified/bulk-rnaseq. The workflow for conducting differential expression analysis is available here: https://github.com/latch-verified/diff-exp. The workflow for conducting functional enrichment analysis is available here: https://github.com/latch-verified/pathway. The LVBRS workflow is built using the Latch Software Development Kit (SDK), which is available as a Python package. Documentation of how to use the SDK for workflow development is available here: https://docs.latch.bio/.

## Authors’ contributions

HGBHL and JLS conducted the analysis. JLS and HGBL wrote the manuscript. NM designed the user interface. MS, AA, AK, RK, and KW developed the LvBRS pipeline. MS, AA, AK, RK, and KW built the Latch infrastructure supporting the workflow orchestration. HGBHL, KG, AA, and KW supervised and coordinated the project. All authors tested the toolkit. All authors read and approved the manuscript.

## Acknowledgments

We thank the Latch AI Inc. team for their helpful discussions and comments. We also thank the Latch software development kit community for their helpful comments.

## Supplementary figures and legends

**Figure S1.**
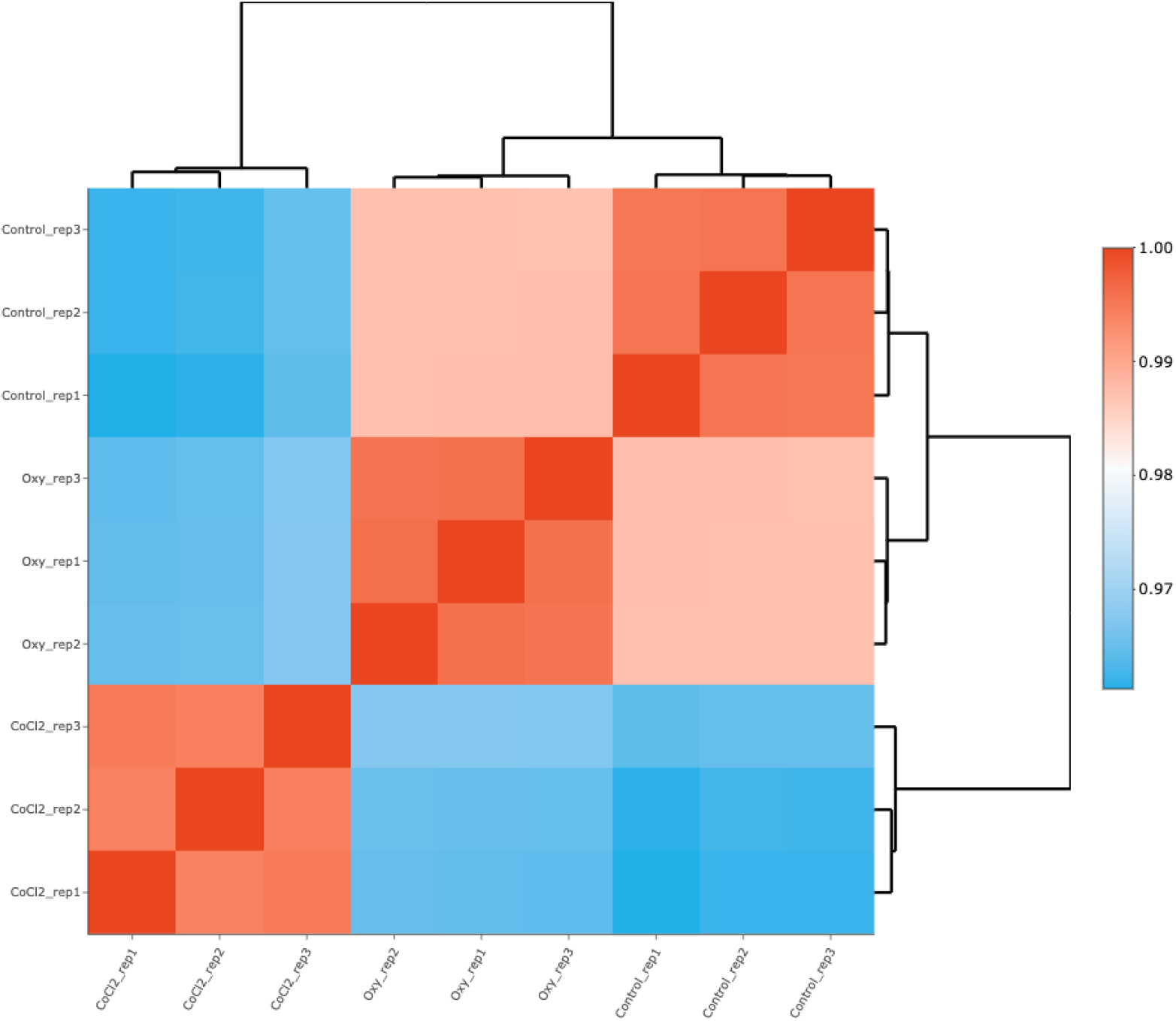
Counts correlation matrix. Samples are hierarchically clustered. Groups with similar global read counts per gene will be closer together; less correlated samples will be further apart. As expected, within-treatment correlations are high, whereas between-treatment correlations are lower. Blue represents lower correlation values; red represents higher correlation values.

**Figure S2.**
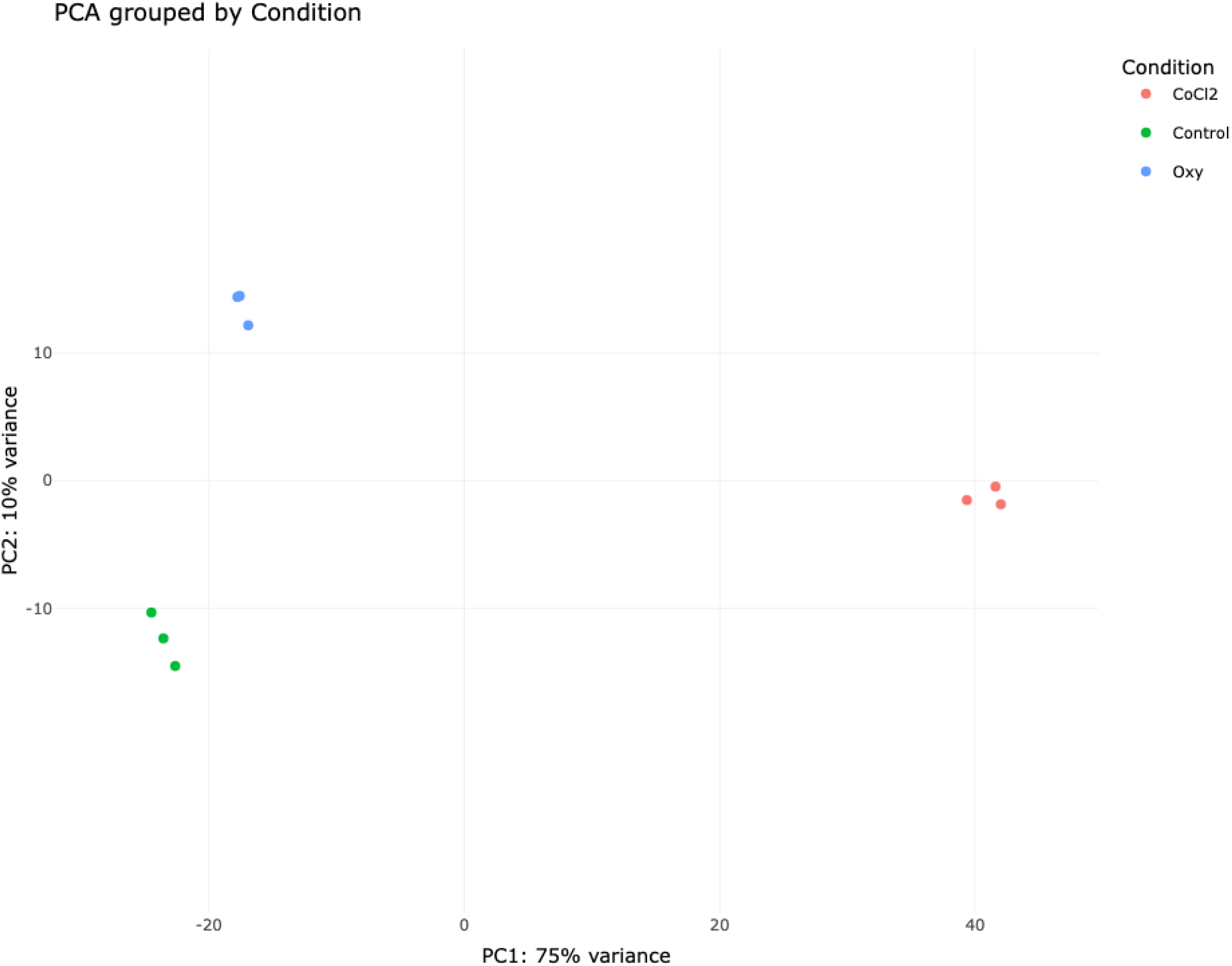
Principal component analysis between samples. Similarities and differences between samples are depicted in principal component space. In this example, 75% of the variance is captured along the first dimension; 10% of variance is capture along the second dimension. Samples from the same condition cluster together.

**Figure S3.**
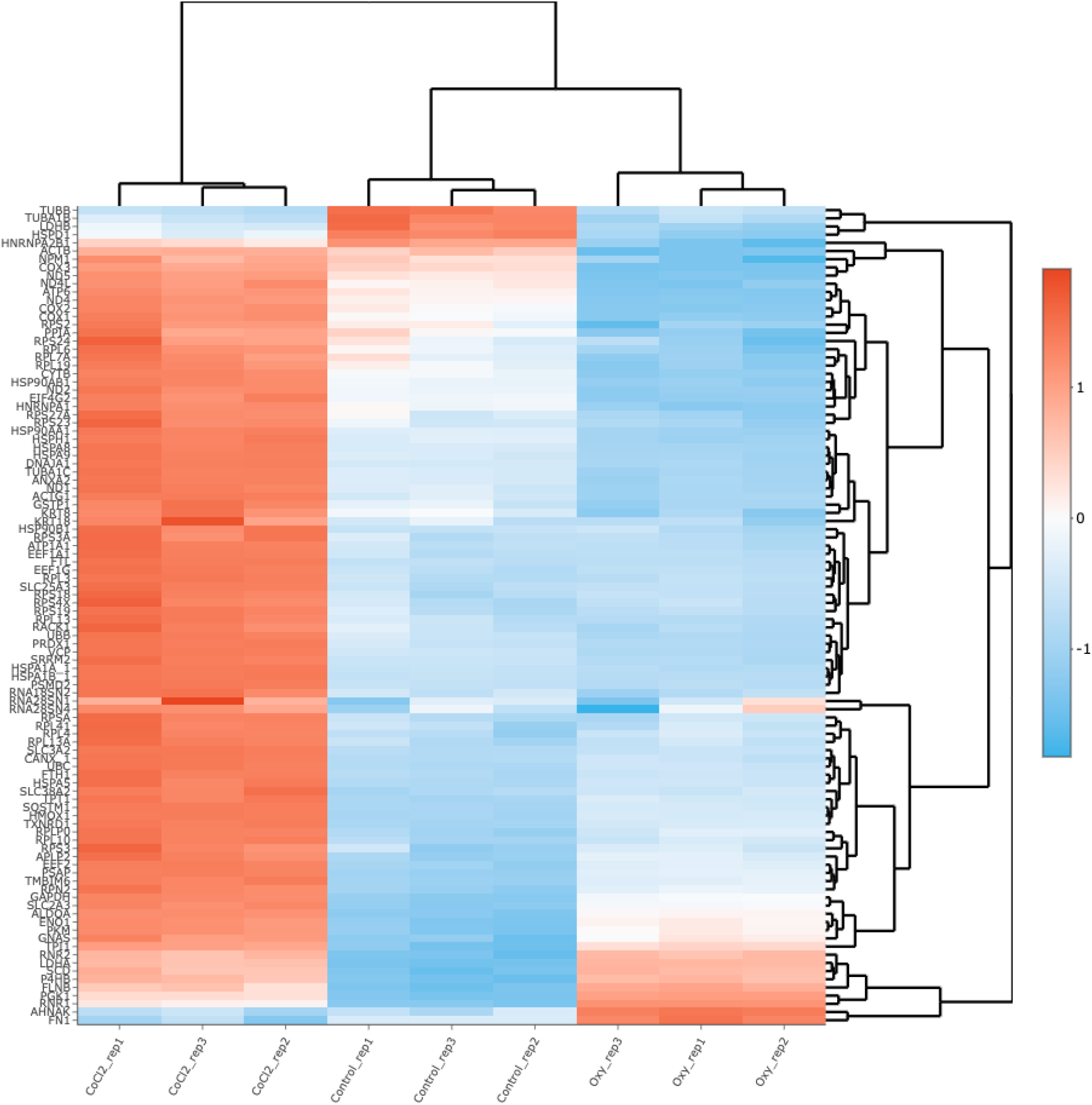
Heatmap of genes with the highest read counts. Initial insights between samples and conditions can be investigated by examining differences in gene expression between samples. Read counts are normalized to z-scores. At most, the top 100 most-expressed genes—defined as having the highest rank of maximal expression across all samples—are depicted. Blue indicates highly expressed genes; red represents lowly expressed genes.

**Figure S4.**
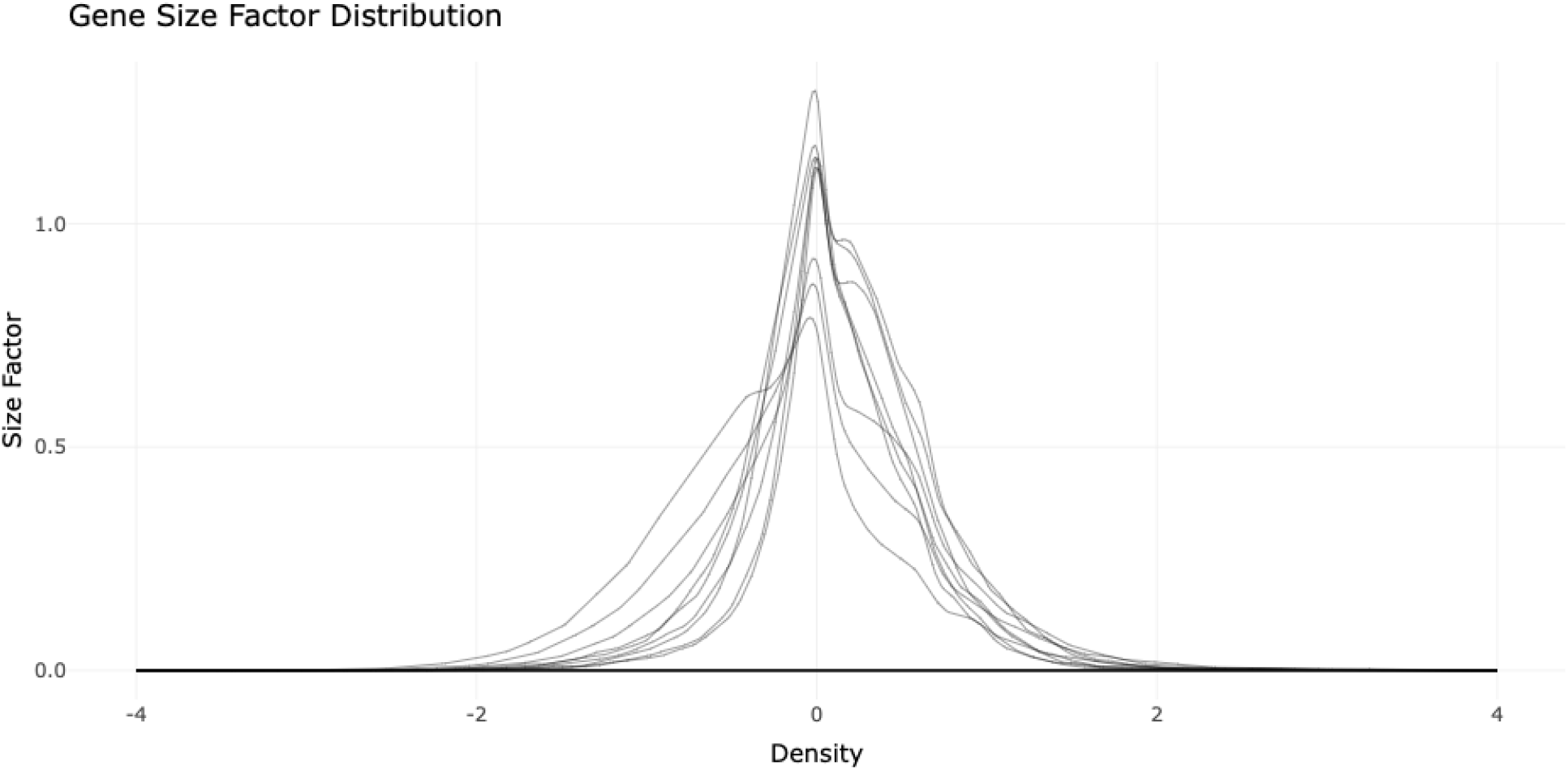
Gene size factor distribution. A tight distribution around zero indicates few genes, which appear in the tails of the distributions, are differentially expressed. Diffuse distributions may indicate large-scale differences among DEG or may warrant further analysis of data quality.

**Figure S5.**
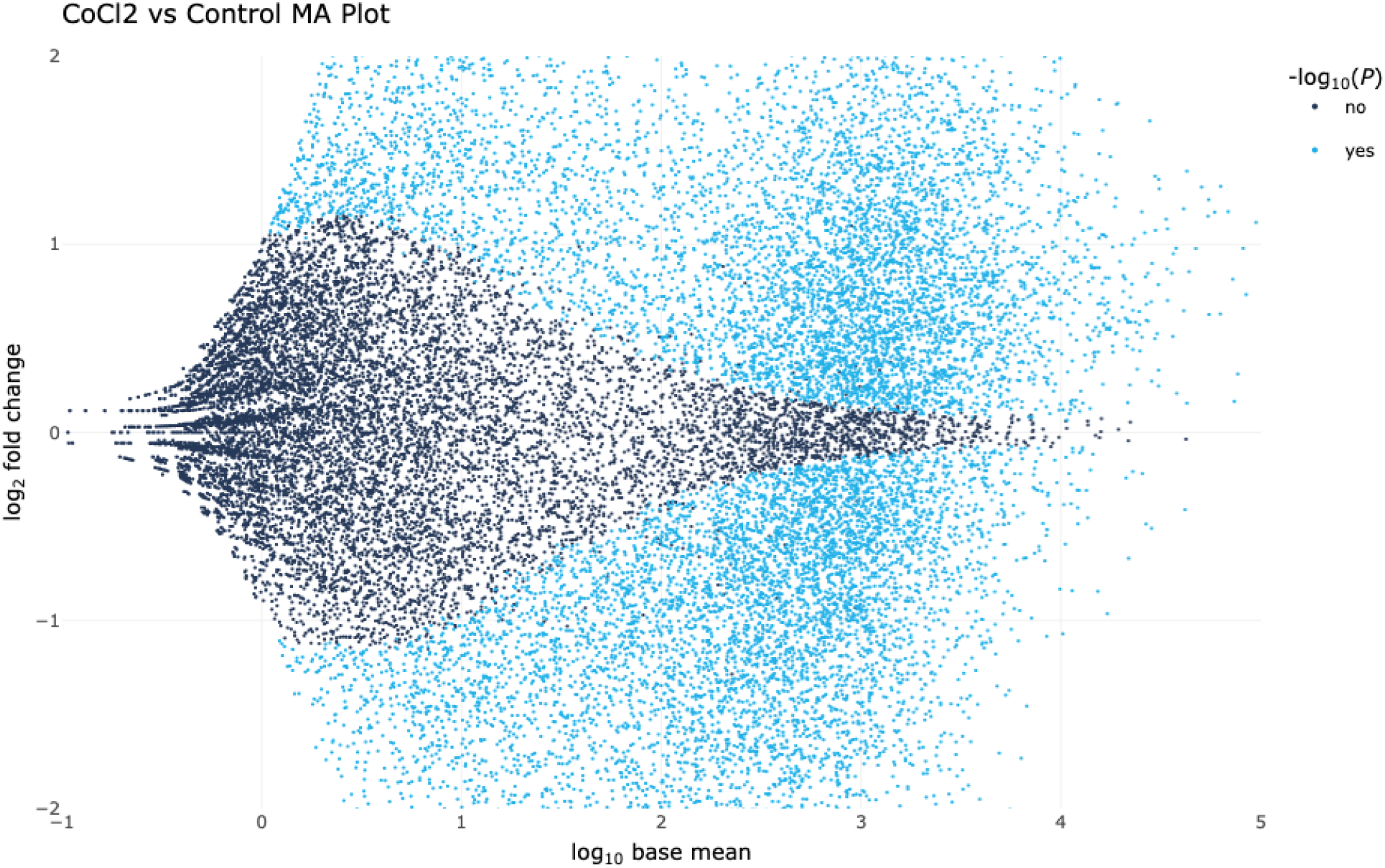
MA plot. A distribution of genes with the largest and smallest fold changes are depicted here, providing a broad overview of how differentially expressed genes are between treatment and control groups. Each dot represents a single gene.

**Figure S6.**
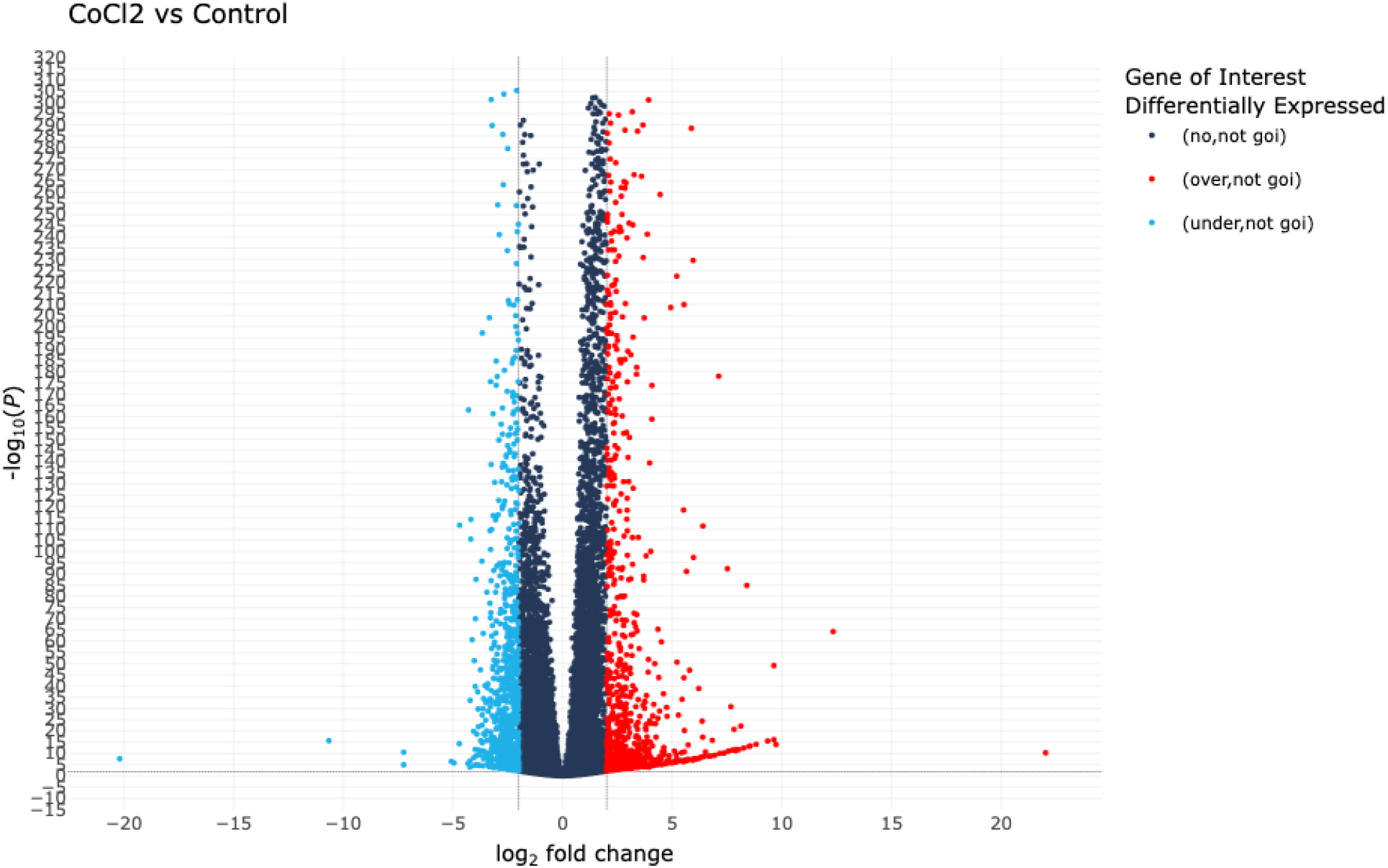
Volcano plot. Differences in gene expression between treatment and control groups are depicted here. The x-axis is the log_2_ fold change; the y-axis is the negative log_10_ p-value. Dots are colored according to if they are not differentially expressed (dark blue), over-expressed (red), or under-expressed (blue).

**Figure S7.**
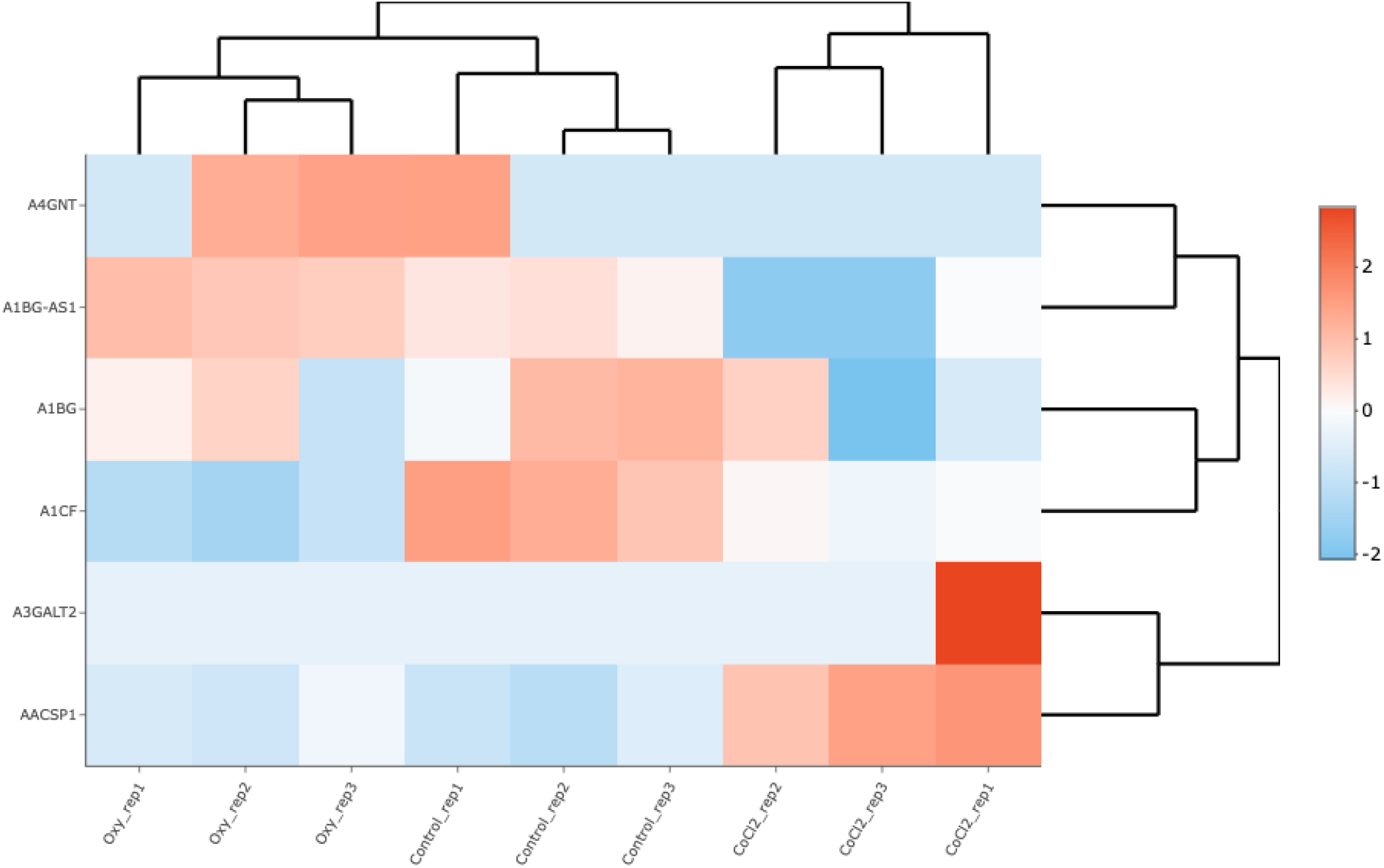
Heatmap of genes of interest. A clustered heatmap of read counts per gene are normalized using a z-transformation. Genes are clustered along the y-axis, and conditions are clustered along the x-axis. Genes that are highly expressed are shown in red; genes that are lowly expressed are shown in blue. Here, a randomly selected six genes—*A4GNT, A1BG-AS1, A1BG, A1CF, A3GALT2*, and *AACSP1*—were chosen for demonstrative purposes.

**Figure S8.**
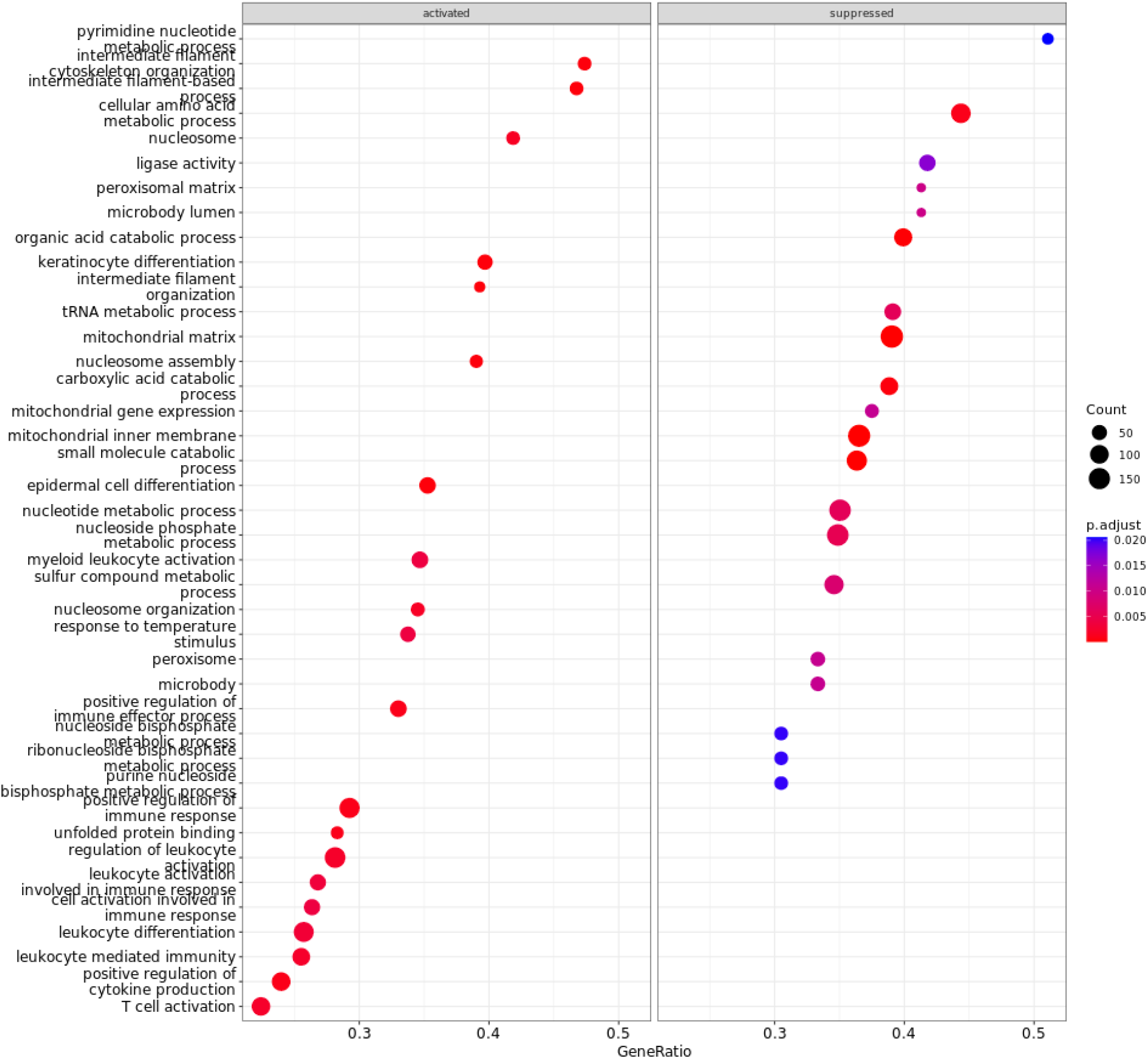
Dotplot of enriched terms. Enriched terms among over- or under-expressed genes are shown on the left or right. Dot size corresponds to the number of genes associated with the term. The color of each dot represents the p-value after multi-test correction. Blue represents less significance; red represents more significance. Depicted here are results using the Gene Ontology database; however, the same figures are generated for the KEGG and MSig databases.

**Figure S9.**
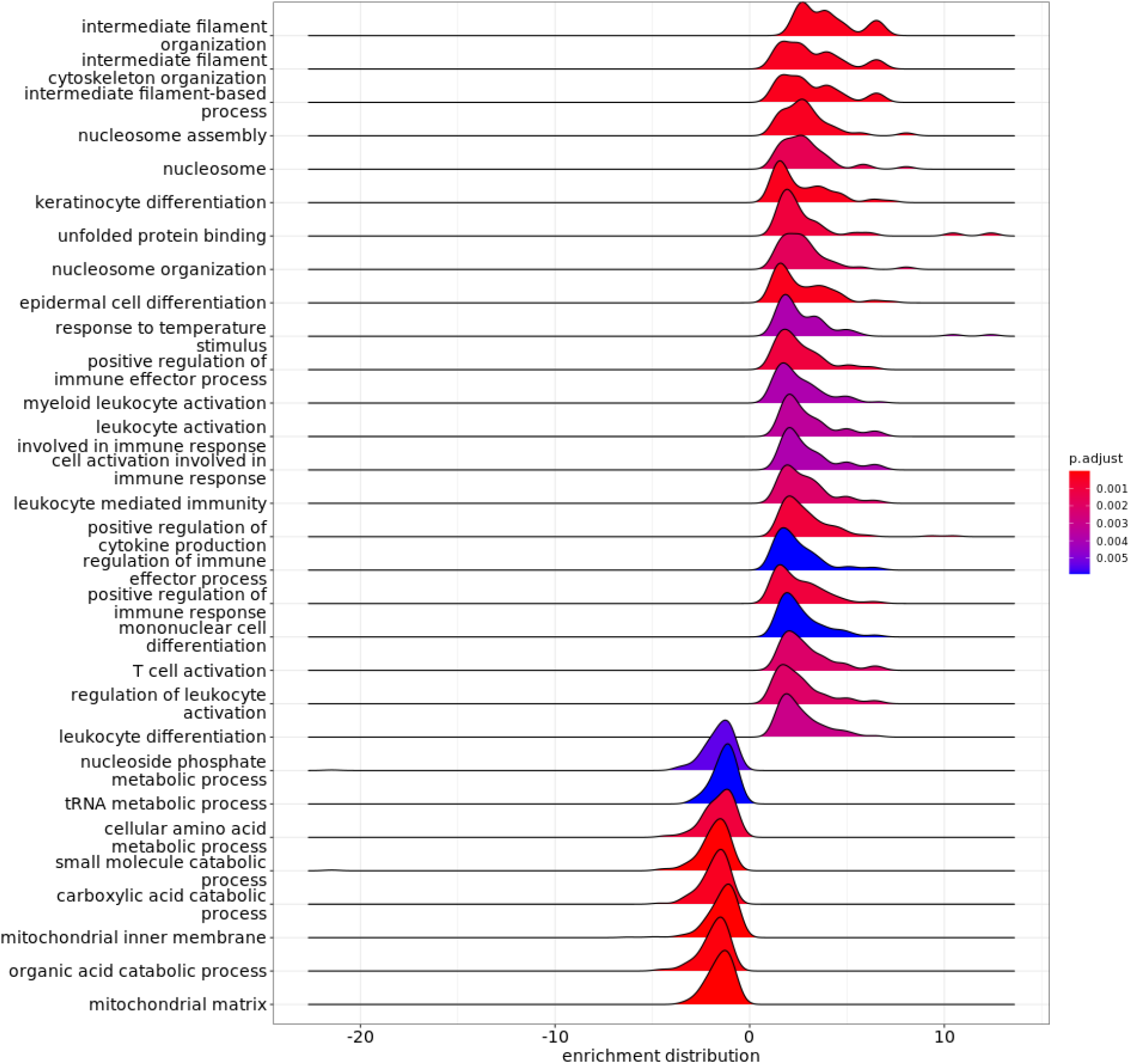
Ridgeplots of enriched terms. Enriched terms and the associated enrichment distributions are depicted here. The color of each distribution represents the p-value after multi-test correction. Blue represents less significant; red represents more significant. Depicted here are results using the Gene Ontology database; however, the same figures are generated for the KEGG and MSig databases.

**Figure S10.**
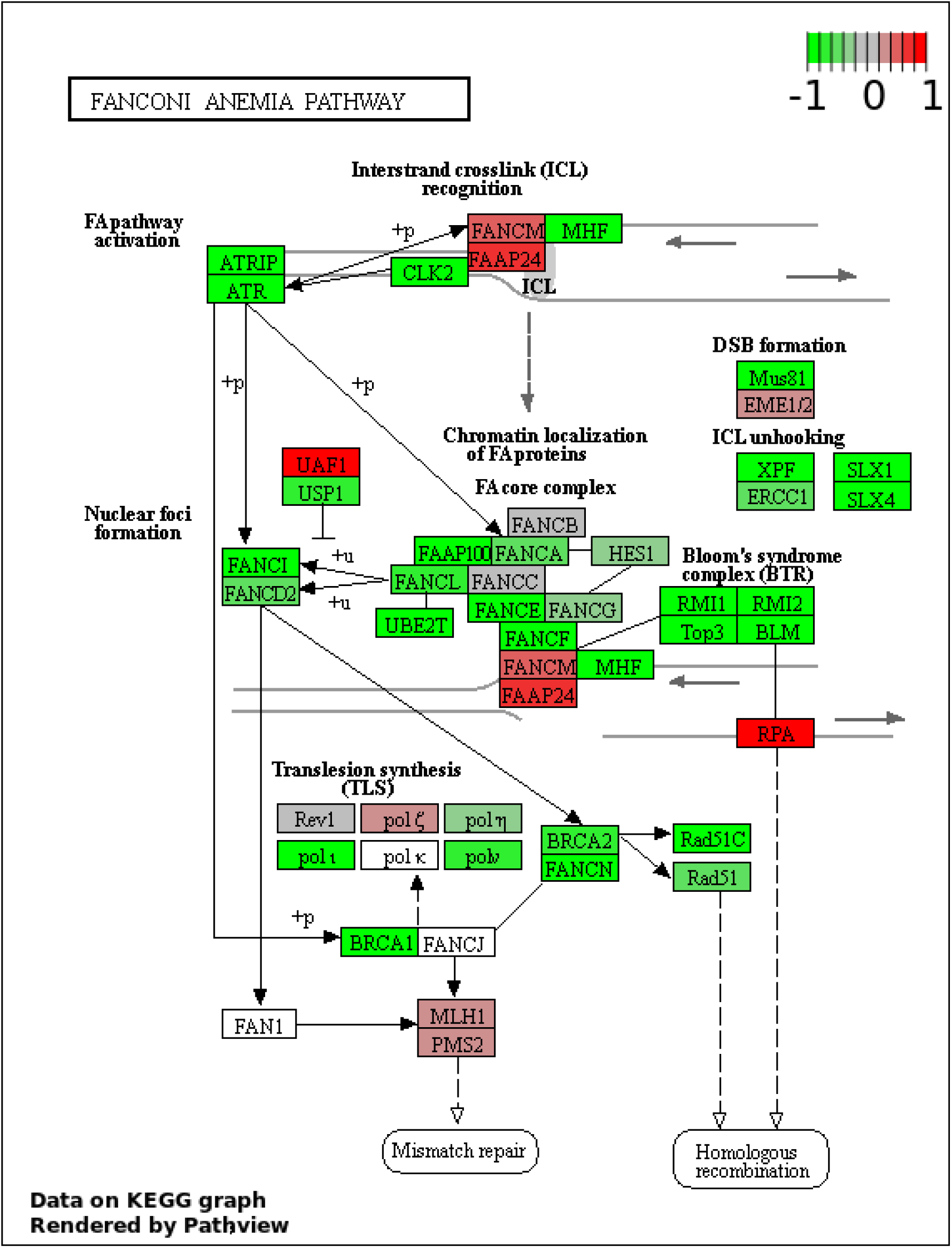
Exemplar KEGG pathway figure. The Fanconi Anemia pathway is depicted as an exemplar KEGG graph, depicting over- and under-expression of genes green and red, respectively. The image is generated using Pathview [27].

**Figure S11.**
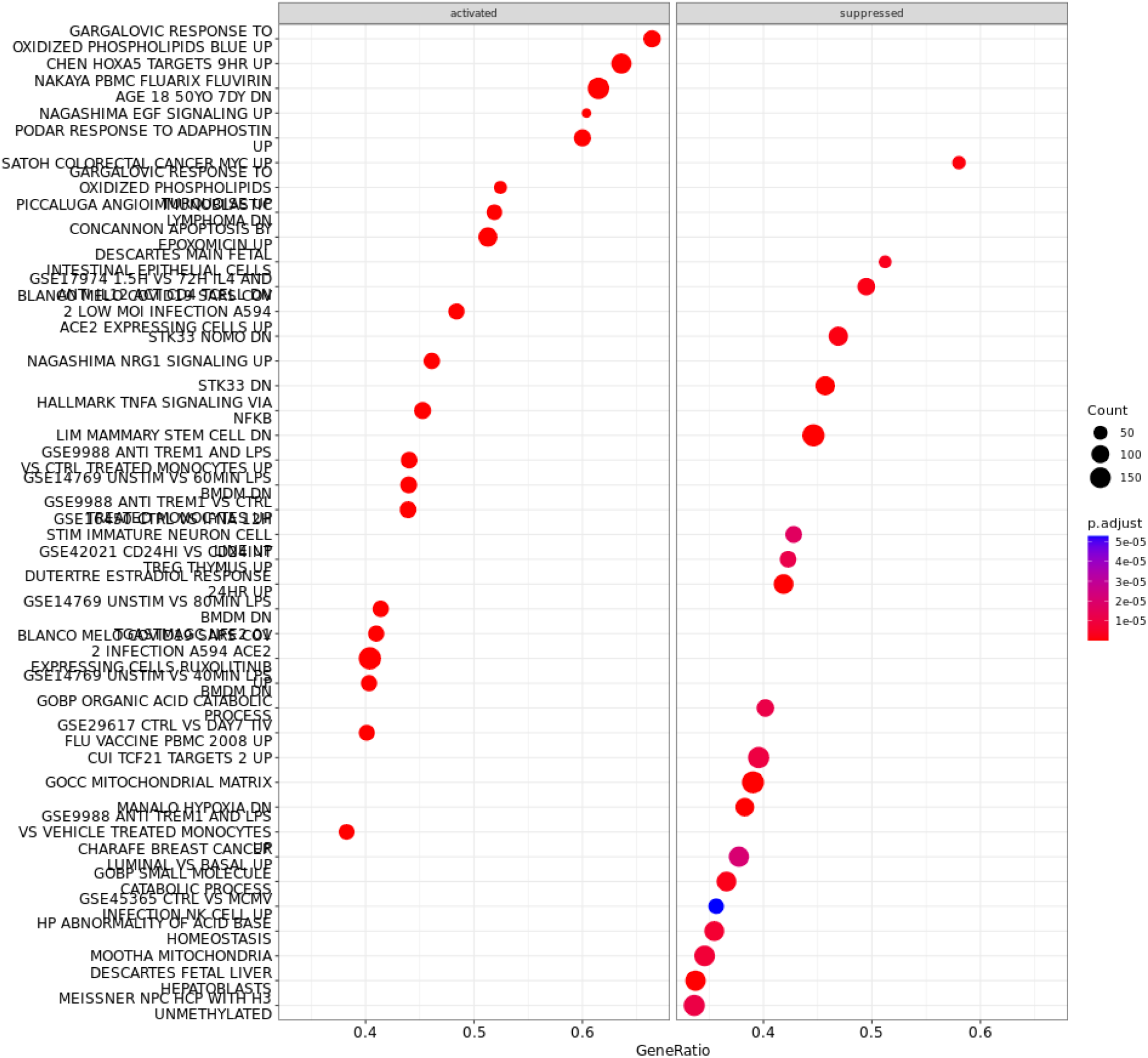
Dotplot of enriched terms using the mSig database. Enriched terms among over- or under-expressed genes are shown on the left or right. Dot size corresponds to the number of genes associated with the term. The color of each dot represents the p-value after multi-test correction. Blue represents less significance; red represents more significance. Depicted here are results using the Gene Ontology database; however, the same figures are generated for the KEGG and MSig databases.

**Figure S12.**
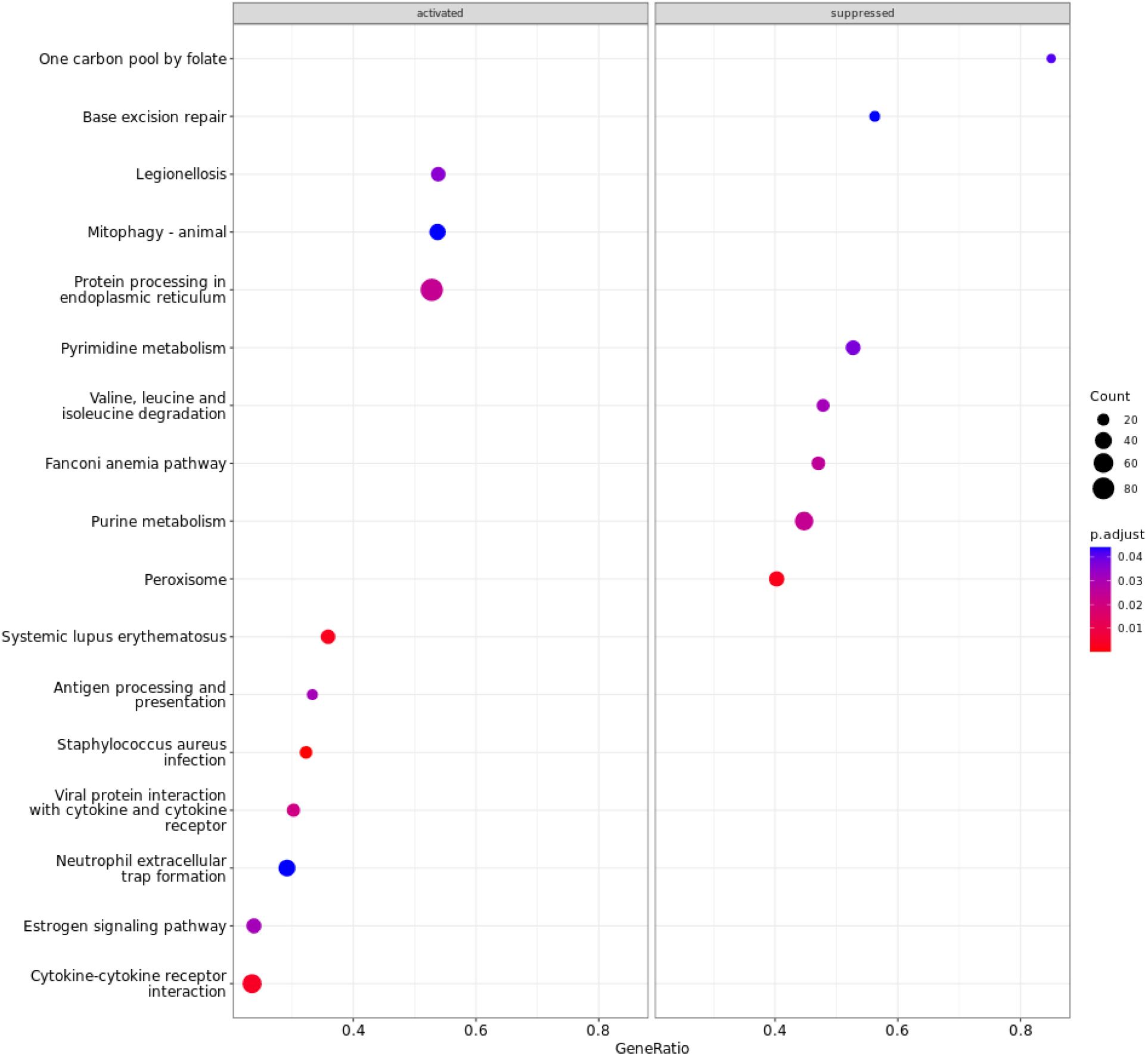
Dotplot of enriched terms using the KEGG database. Enriched terms among over- or under-expressed genes are shown on the left or right, respectively. Dot size corresponds to the number of genes associated with the term. The color of each dot represents the p-value after multi-test correction. Blue represents less significant; red represents more significant. Depicted here are results using the Gene Ontology database; however, the same figures are generated for the KEGG and MSig databases.

